# Addiction beyond pharmacological effects: the role of environment complexity and bounded rationality

**DOI:** 10.1101/179739

**Authors:** Dimitri Ognibene, Vincenzo G. Fiore, Xiaosi Gu

## Abstract

Several decision-making vulnerabilities have been identified as underlying causes for addictive behaviours, or the repeated execution of stereotyped actions despite their adverse consequences. These vulnerabilities are mostly associated with brain alterations caused by the consumption of substances of abuse. However, addiction can also happen in the absence of a pharmacological component, such as seen in pathological gambling and videogaming. We use a new reinforcement learning model to highlight a previously neglected vulnerability that we suggest interacts with those already identified, whilst playing a prominent role in non-pharmacological forms of addiction. Specifically, we show that a dual-learning system (i.e. combining model-based and model-free) can be vulnerable to highly rewarding, but suboptimal actions, that are followed by a complex ramification of stochastic adverse effects. This phenomenon is caused by the overload of the capabilities of an agent, as time and cognitive resources required for exploration, deliberation, situation recognition, and habit formation, all increase as a function of the depth and richness of detail of an environment. Furthermore, the cognitive overload can be aggravated due to alterations (e.g. caused by stress) in the bounded rationality, i.e. the limited amount of resources available for the model-based component, in turn increasing the agent’s chances to develop or maintain addictive behaviours. Our study demonstrates that, independent of drug consumption, addictive behaviours can arise in the interaction between the environmental complexity and the biologically finite resources available to explore and represent it.

## Introduction

Addiction is marked by the compulsive execution of stereotyped actions despite their adverse consequences [1, 2, 3, 4, 5]. This maladaptive form of decision making is typically associated with the consumption of substances of abuse, such as alcohol, tobacco, illicit and prescription drugs [3, 6, 7]. More recently, the definition has been also used to describe gambling [8, 9] and other putative forms of behavioural addictions, such as internet gaming [10]. Importantly, these latter forms of addiction lack the neuro-pharmacological effects of a consumed drug, and yet are characterised by a striking similar symptomatology.

Several theories and computational models have been proposed to explain the repetition of suboptimal decisions typical of addiction [9, 3, 6, 7]. These theories assume decision making results from the interaction of multiple systems, e.g. habitual, deliberative, Pavlovian, motivational, situation identification, etc. which rely on different learning and computing principles. This composed structure is associated with several vulnerabilities to the pharmacological effects of drugs of abuse, each of which can result in the expression of compulsive repetition of drug intake [3, 11, 12, 13, 14, 15].

In particular, Reinforcement Learning (RL) models of addiction frequently assume that aberrant drug-seeking habits come to dominate behaviour in addiction due to drug induced bio-chemical hijacking of the dopaminergic prediction error signal [7, 16, 3, 17, 18, 19, 20]. The hypothesis of the dominance of the habitual system nicely accounts for aspects of addiction such as inelastic behaviour in the face of changes in the environment or even in presence of punishing outcomes following drug consumption [16, 21]. However, several other behaviours associated with addiction are left unaccounted for [18, 3]. First, one of the defining characteristics of substance abuse according to the DSM-5 is “A great deal of time is spent in activities necessary to obtain the substance (e.g., visiting multiple doctors or driving long distances)” [22]. Such temporally extended activities are often novel, complex and context-dependent [23, 18, 3, 24], and therefore are not driven by habitual processes or stimulus-response conditioning. Second, phenomena such as craving can occur even without exposure to conditioned stimuli (but see [25, 26]). Finally, gambling [8, 2, 1] and internet gaming [10], which are also considered part of the addictive behaviours, lack the pharmacological interference that is considered essential to drive the aberrant habit formation [9].

These issues have been partially addressed by hypothesising the presence of vulnerabilities affecting the deliberative system [3]. In particular, it has been suggested that non-habitual forms of addictive behaviours may be caused by errors of interpretation, where either the outcome of an action (drug consumption, gambling etc.) is over-evaluated as beneficial or useful or the long term consequences of these actions are under-evaluated in their negative effects. However, the computational mechanisms by which both drug-related and non drug-related addiction can induce these effects on the deliberative/planning system are not well understood [11, 27, 28].

Other models [9] have posed that addiction can emerge in environments characterised by incomplete or inaccessible information. Under these conditions, the underestimation of the negative consequences or the over-evaluation of the positive ones is simply caused by a lack of information. However, this hypothesis does not seem to match with clinical evidence, as once the required information is made readily available to addicted individuals, motivating their abstinence, relapse should not occur.

We propose a solution can be found in the analysis of the discrepancy between the resources available to an agent and those required to explore, represent or compute the environment it operates in. Most computational models of addiction have so far focused on environments characterised by the presence of easy to compute outcomes, where the number of actions available and their ramifications were limited. This simplification has distanced the computational analysis from the clinical practice, which has long considered a wide range of environmental factors, and social interactions in particular, to have a strong impact on addiction development and maintenance [29, 30, 31].

Environment complexity and exploration are well recognised factors in the fields of Artificial Intelligence (AI) [32, 33], as well as developmental and computational neuroscience, in particular when considering the problem of the exploration-exploitation trade-off [34, 35, 36, 37, 38]. As the amount of experience required by an agent to achieve a specific behavioural performance grows faster than the product of the number of available states and actions [39, Chapter 8], exploration and training in complex environments can easily result in incomplete or incorrect representations of action-outcome ramifications [37, 40, 41]. Furthermore, if a complex environment is correctly represented in the agent’s internal model, e.g. after a prolonged exploration, the stored action-outcome ramifications might still overload the agent’s capacity to *internally assess* its available options. This inherent inadequacy of resources can be also aggravated by temporary forms of cognitive impairments which would dynamically increase the chances to trigger suboptimal planning. Interestingly, anxiety or stress are good examples of dynamic processes associated with temporary cognitive impairments and represent known triggers in addiction disorders and relapse after treatment [42, 43, 44, 45].

Our simulations show that the development of addictive behaviours may be supported by the interaction between specific features of the environment and both habitual and deliberative processes [37, 40, 46, 47]. We propose this vulnerability complements and interacts with previously described ones, capturing the emergence of addiction in the absence of pharmacological factors.

## Materials and Methods

### Agent

The behaviour of our simulated agents (Fig 1) is controlled by a hybrid (or dual) RL model system [48, 18, 49, 50, 51]. This algorithm maximises expected cumulative rewards by simultaneously learning, through a model-free (MF) component, and computing, through a model-based (MB) component, an optimal action strategy, or policy *π*.

**Figure 1:**
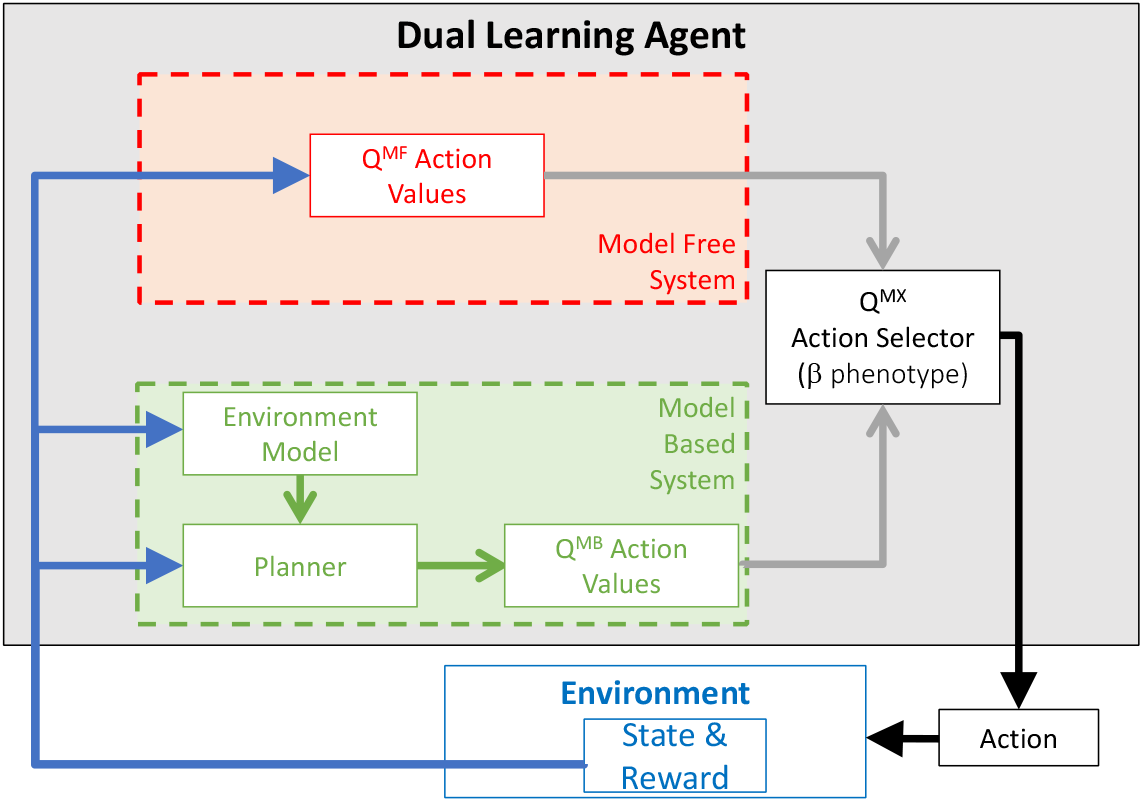
Dual Learning Agent. The decision making architecture includes: (i) a model free component (MF), which updates action values using value prediction error computations; and (ii) a model based component (MB), which generates an internal model of the environment, based on experienced action-outcomes and bounded computations. Act ion-out come estimations derived from the two components are combined linearly according to a balance parameter, *β*, to drive action selection.

The MF component is implemented as a standard tabular Q-learning algorithm [52]. MF algorithms such as Q-Learning and actor-critic architectures [53] are usually employed to model habitual behaviours [54, 55, 3, 48, 56, 57] and therefore are tipically associated with the dorsal cortico-striatal neural circuit [49, 58]. These algorithms are characterised by limited flexibility but computational efficiency as they require limited resources to slowly update associations between state-action pairs and values 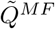(*s, a*), depending on experience. The MB algorithm is employed to implement planning processes [11, 27, 26, 41] on the basis of an explicit representation, in an *internal model*, of action-state relationships and associated rewards, as experienced in the environment. Due to the similarity with goal-oriented processes, the MB component is often associated with the ventral cortico-striatal circuit [49, 58]. Where the MF component simply selects the best action among those available in its current state, the internal model of action-state sequences allows the MB component to evaluate entire policies, as if navigating decision trees with their ramifications and consequences, before making any decision. Such a process of evaluation is demanding in terms of computational resources and time, but allows a high degree of flexibility.

Most dual models assume an ideal MB process [50, 59], characterised by a complete knowledge of the environment and unlimited computational resources, which therefore always leads towards optimal choices. However, biological MB system are constrained, or *bounded,* by their limited resources [60, 61, 62, 63, 64, 65, 66, 67]. Thus, to model biologically plausible healthy and dysfunctional behaviours (as e.g. in addiction [18, 3]), in our simulations we have employed a MB component that represents only direct experience, and that relies on bounded computational resources [60, 61] to navigate its internal model. Importantly, our MB component generates a new value estimation at each step by applying the Bellman Equation a limited number of times to states sampled stochastically, following an early-interrupted variation of the Prioritized Sweeping algorithm [68], with stochastic selection of the states to update (see Algorithm 1). This is similar to what is regularly done in the Monte-Carlo Tree Search family of algorithms [69], which is commonly adopted in Artificial Intelligence for complex environments models where estimations over simulations are easier than complete bellman backups. However, the Early Interrupted Stochastic Prioritized Sweeping algorithm employed here is computationally more efficient for small environments [70], so to provide stable results with a limited number of updates.

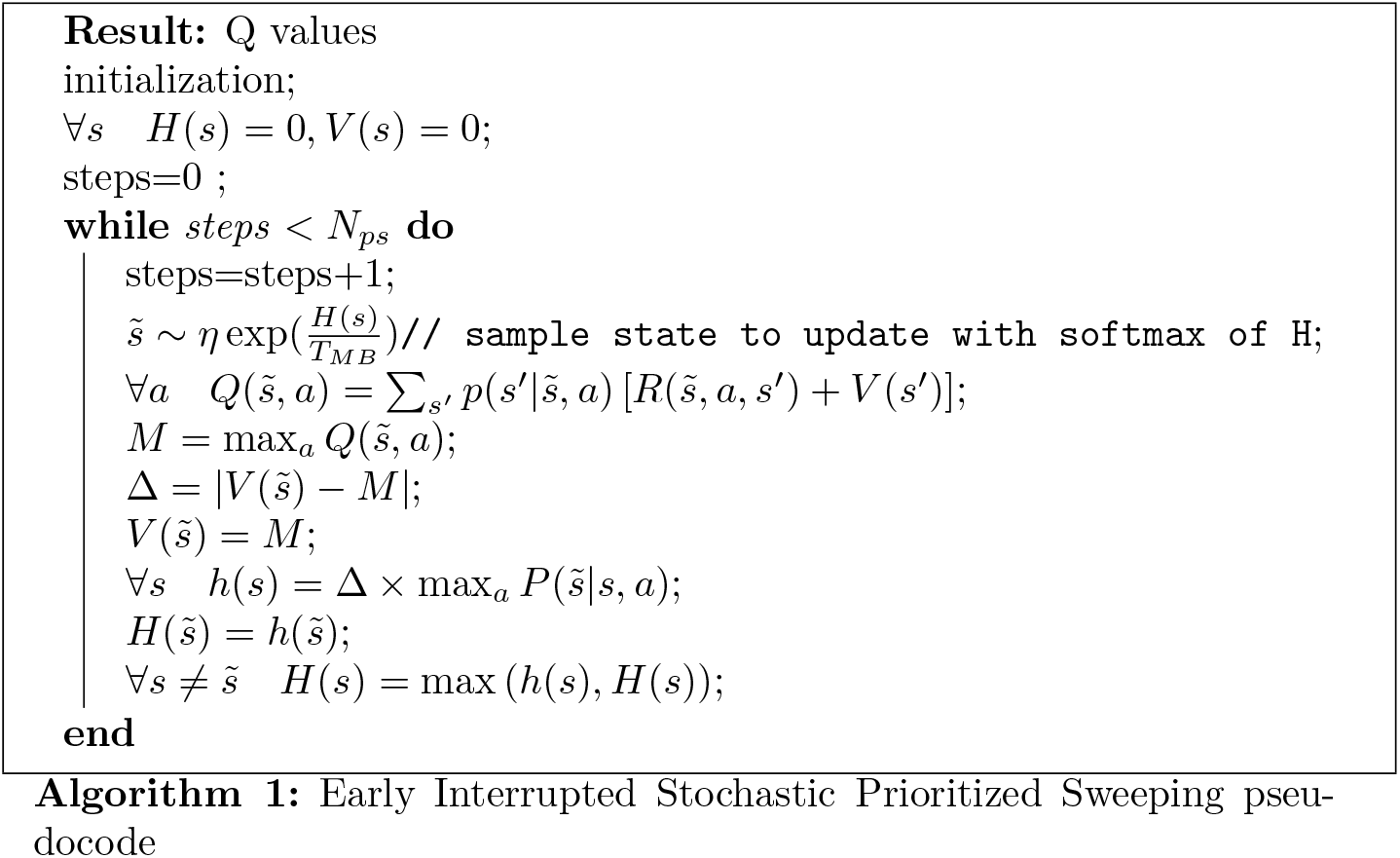

In keeping with existing literature [11], we assumed that the MB and MF components do not share a common representations, and they do not interact during the computation of the respective state action values. However, a hybrid value function *Q^MX^* is computed by balancing MF (*Q^MF^*(*s, a*)) and MB (*Q^MB^*) estimates depending on a parameter, *β*, as follows:

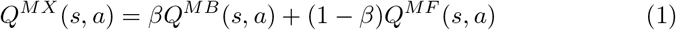

Similar to a previous study [58], six values (1, 0.8, 0.6, 0.4, 0.2, 0) have been used for this parameter to simulate different behavioural phenotypes, along a spectrum between purely model-based (*β*=1) and purely model-free (*β*=0) reinforcement learning. In terms of neural implementation, these phenotypes loosely match the neural systems dominated by either a ventral or a dorsal cortico-striatal circuit, with the strength of the directed connectivity between these circuits as the analogue of the beta values in the algorithmic model.

Finally, the agents selected the actions that were expected to maximize the future utility (*Q^MX^*) in 90% of their selections. For the remaining 10% of selections, the agents would perform a random action, in a standard strategy meant to preserve exploration for all stages of the simulations, termed stochastic e-greedy *persistent exploration* [71].

### Environment

We tested our hypothesis that suboptimal, addiction-like, behaviours can emerge without pharmacological interference or MB-MF malfunction, in an environment (Fig 2) that allows long action-sequences characterised by deep ramifications. In comparison with simpler environments, characterised by limited interactions or depth of action sequences (e.g. an operant conditioning chamber), environments simulating open space navigations require larger amount of resources invested in the exploration and computation of the action-outcome contingencies. Thus, the agents struggle to find and pursue those policies that lead to reward maximization (i.e. optimal behaviours) and to avoid those policies that lead overall losses (i.e. suboptimal behaviours).

**Figure 2:**
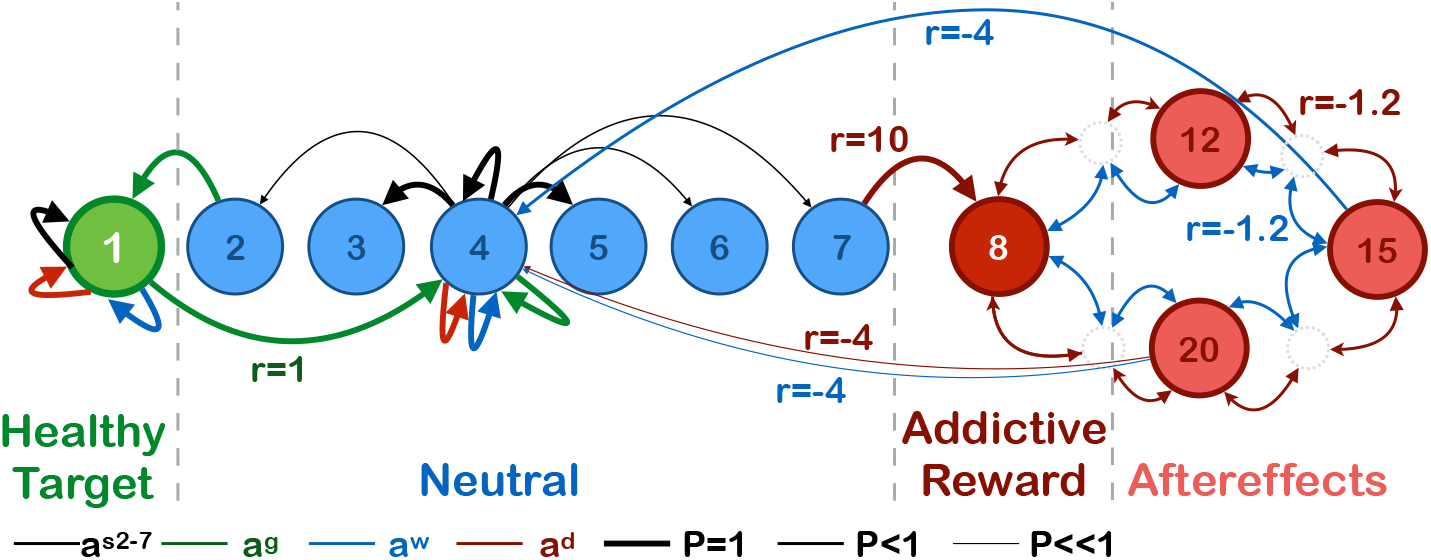
Illustrative representation of the environment. The states are disposed in a linear arrangement: on the left (number 1) a state associated with a healthy reward, on the right (number 8) a state associated with an addictive non-drug reward (e.g. gambling), separated by 6 neutral states that can be freely traversed. Entering the healthy reward state results in a moderate reward (*R_g_* = 1), after which the agent returns to the central neutral state (number 4). Entering the addictive state provides an immediate high reward (*R_g_* = 10), followed by a further segment of 14 states that are associated with negative outcomes (−1.2) or punishments. Within this segment of *after-effects*, action results are stochastic, making it difficult for the agent to find a way out of this part of the environment, and resulting in an average overall punishment that makes the selection of the addictive reward suboptimal. In this illustrative representation, few key transitions are reported, with detailed descriptions for the states 1,4,15 and 20 for which line width represents transition probabilities and colour represents the action class (*a_s_, a_g_, a_w_, a_d_*). Neutral states can be crossed by selecting actions *a_s_*2–7, which are deterministic for adjacent state while have high chance of failing for distant states. Agents can reach the healthy reward state by executing action αg whilst in state 2, and the addictive reward state, by executing action *a_w_* whilst in state 7. In the after-effect segment, actions results are less predictable and only action *a_w_* at state 15 has a high chance of leaving the addictive area, with a cost of −4. All details about the environment are reported in table 2.

Importantly, we could not investigate the same phenomena by including, for instance, a high discount factor in a simplified environment, as there are fundamental differences between disregarding temporally distant events and failing in exploring, representing and evaluating them. In fact, with a high discount, an addictive behaviour that disregards long term negative effects would be formally optimal and therefore it would not induce that sense of inability to stop [19] that often characterizes addiction.

The simulated agents operated under two different configurations of the environment or phases (Fig 2). Under the initial *safe phase* (*d_init_* = 50 steps), the agents could only experience a moderate reward (termed *healthy* reward, *R_g_* = 1) if they accessed the relative state. Once the healthy reward state was reached, an agent would be brought back to the initial state and could pursue the reward again. No other reward or punishment was available in any other part of the environment. Under the second *addiction phase* (*d_drug_* = 1000 steps) the agent was still rewarded by accessing the healthy reward state, but it could also access a state characterised by a high reward (termed *addictive* reward, *R_d_* = 10). This state was inescapably followed by a more unpredictable and mixed-in-value negative *after-effect segment* of the environment, which ideally simulated the multifaceted effects addictive behaviour has on the social life and health of the addicted individual. At the end of this after-effect segment, the agent would be again brought back to the initial state. Table 1 shows the number of updates that the original Prioritized Sweeping algorithm would have used to find the optimal policy in each phase. These are two orders of magnitude larger than the updates allowed by the adopted bounded MB.

**Table 1:** Number of updates necessary to Prioritized Sweeping to find the value function for each phase

| Phase | Number of Updates |
| --- | --- |
| Init | 4,712 |
| Addictive Reward | 5,005 |

**Table 2:** Environment Model Parameters

| Name | Description | Value |
| --- | --- | --- |
| $N_T$ | Number of states | 22 |
| $N_G$ | Number Goal States | 1 |
| $N_D$ | Number Addictive Area States | 15 |
| $N_n$ | Number Neutral States | 6 |
| $N_a$ | Number of actions | 9 |
| $S_0$ | Starting state | 4 |
| $R_p$ | Punishment end of Addictive Area | -4 |
| $R_c$ | Punishment in Addictive Area | -1.2 |
| $R_{dd}$ | Reward at entering Addictive reward state | 10 |
| $R_g$ | Reward when entering healthy reward state | 1 |
| $d_{init}$ | Duration safe phase | 50 |
| $d_{drug1}$ | Duration addictive phase | 1000 |

**Table 3:** Agent Model Parameters

| Name | Description | Value |
| --- | --- | --- |
| $\alpha$ | MF learning factor | 0.05 |
| $\gamma$ | Discount factor | 0.9 |
| $d_{MB}$ | MB decay factor | 0.01 |
| MBUS | Number of MB updates | 50 |
| $T_{MB}$ | Temperature for stochastic state update selection | 1 |
| $\epsilon$ | Exploration Factor | 0.1 |

Finally, to test the ability of the agents to adapt to changes, we modified the environment structure in a separate set of simulations. This modified environment included three arms in a Y shape, adding a segment to the two already described. This third segment-termed *neutral*-was kept empty, and reaching its end did not send the agent back to the starting position (as for the healthy reward state) or have it enter an after-effect segment (as for the addictive reward state), but it allowed the agent to freely move to the adjacent neutral states. After the time step 2500, the healthy reward (and its associated rule of sending back the agent to the origin point) was moved from its initial position to the end of the neutral segment. At the same time, the healthy reward segment became neutral (i.e. deprived of any reward), also inheriting the rule of free state transitions among neutral states instead of leading back to the initial state.

## Results

Independent of differences in the parametrisations regulating MB/MF balance, agents seem to rapidly acquire a stable behaviour, marked by the nearexclusive preference for either the healthy or the addictive state (Fig 3). This bifurcation into either an optimal (healthy) or a suboptimal (addictive) be-haviour trajectories is determined by few initial choices. The *healthy behaviour* is reached after less than 300 steps, across populations, and it is maintained for the entire time-length of the experiment. Conversely, the *addiction trajectory* is characterised by long-lasting, albeit transient, choice preferences, which are reached after less than 100 steps. Long simulations employing agents controlled uniquely by the MF component have proven the length of this transient stability is significant. These agents converge to optimum after around 100k steps (Fig 4), in comparison with the 300 steps required by the healthy agents, with identical parametrization, to engage in the optimal behaviour (cf. [52]). It must be noted that the MF component is a standard Q-Learning agent which has been formally proved to converge and which can be easily used to reproduce previous findings related to addiction, once the algorithm is used in association with easy to explore and compute environments [16].

**Figure 3:**
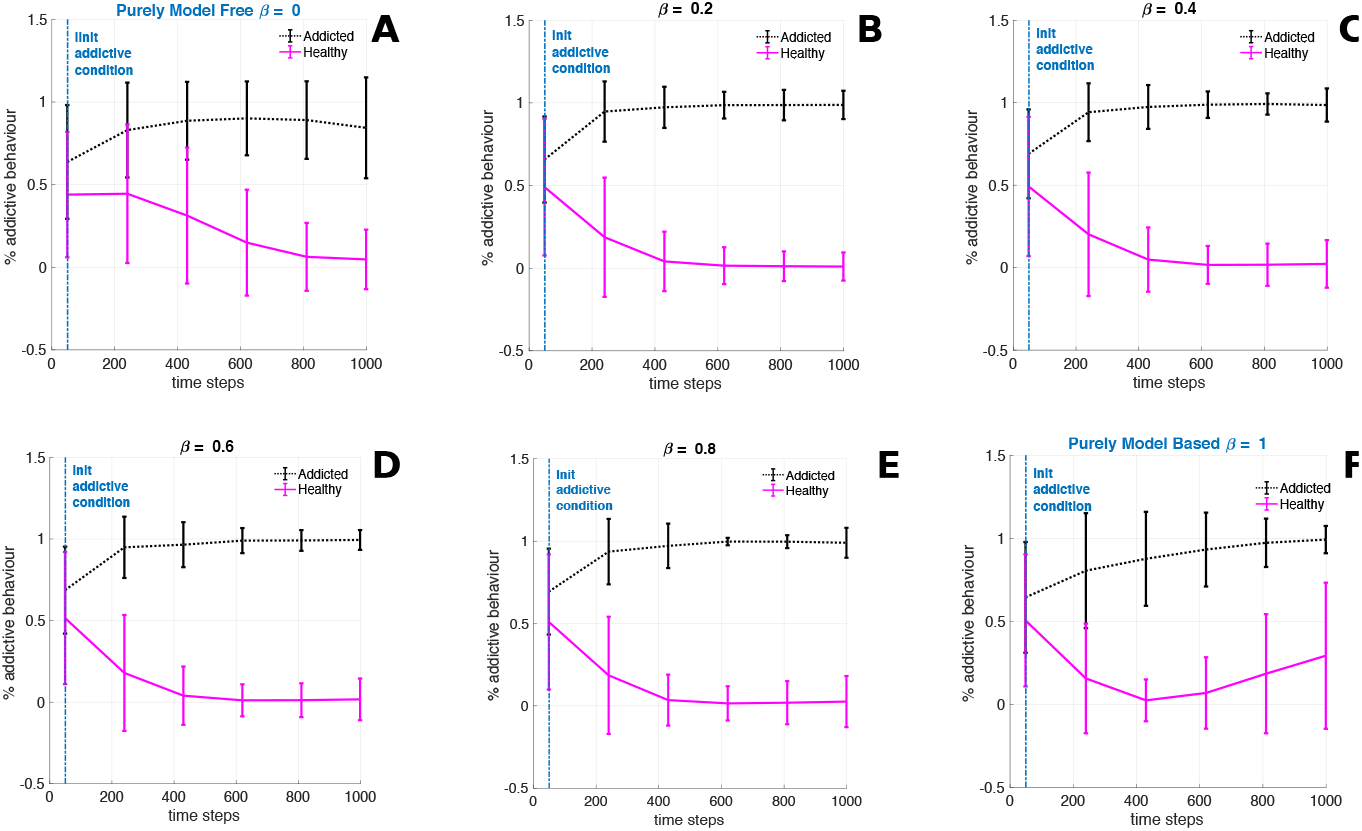
Behavioural trajectories illustrating the ratio of healthy vs addictive reward state selections for addicted and healthy subjects. The six panels highlight different behavioural trajectories, depending on *β* values, which represent MB/MF balance per population. Addicted agents are defined as those visiting the addictive reward state (number 8) more often than the healthy reward state for the whole experiment duration (0:1000 steps). Healthy subjects are defined by subtraction. Each of the six configurations of *β* values was tested with a total of 900 agents (healthy+addicted). Each data point in the chart reports mean and standard deviation for the number of visits to either the addictive of the healthy reward state, over the sum of the total visits to either state, across the 900 agents. A bifurcation in choice preferences clearly emerges between addicted and healthy agents, for all parametrizations.

**Figure 4:**
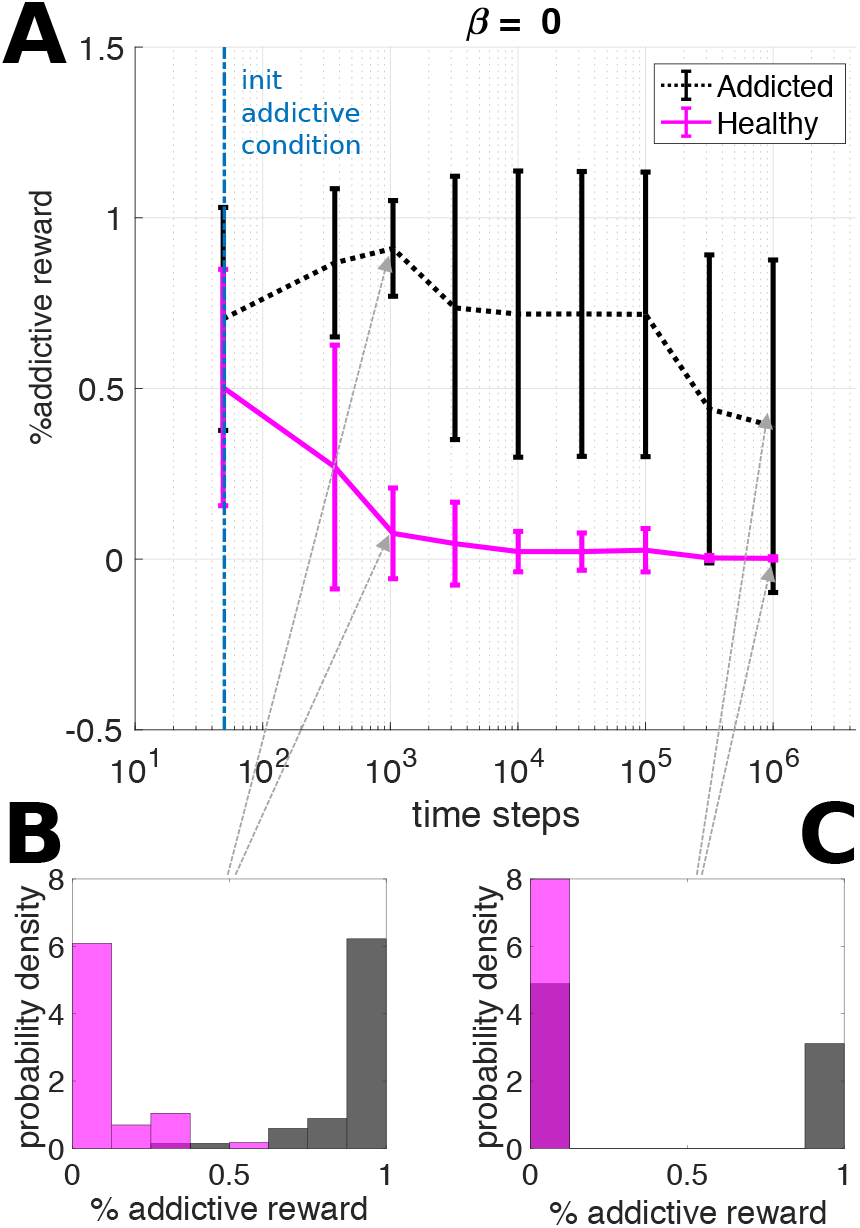
Long runs with logarithmic time scale. Behaviour expressed by purely MF agents (*β* = 0) was recorded and averaged over 100 runs, separating the addicted agents (high preference for the addictive reward state in the first 10,000 steps) from the healthy ones (the remaining agents, which showed the opposite preference within the same time period). A clear bifurcation emerged in the behaviour of the agents (cf. panel A with Fig. 3). Most of the addicted subjects changed their policy towards a healthy behaviour within a time of 200K steps. Histograms in panels B and C also illustrate the behavioural bifurcation, as the behaviour falls either in the interval with the lowest drug intake preference (0-0.125) or in the interval of the highest intake (0.875-1).

In a previous study (cf. [58]), we demonstrated across algorithmic and neural implementation that the balance between MB and MF components significantly affected the chances to develop addictive behaviours, as higher resistance to addiction was found in populations characterised by *intermediate* values of *β* (Fig 5).

**Figure 5:**
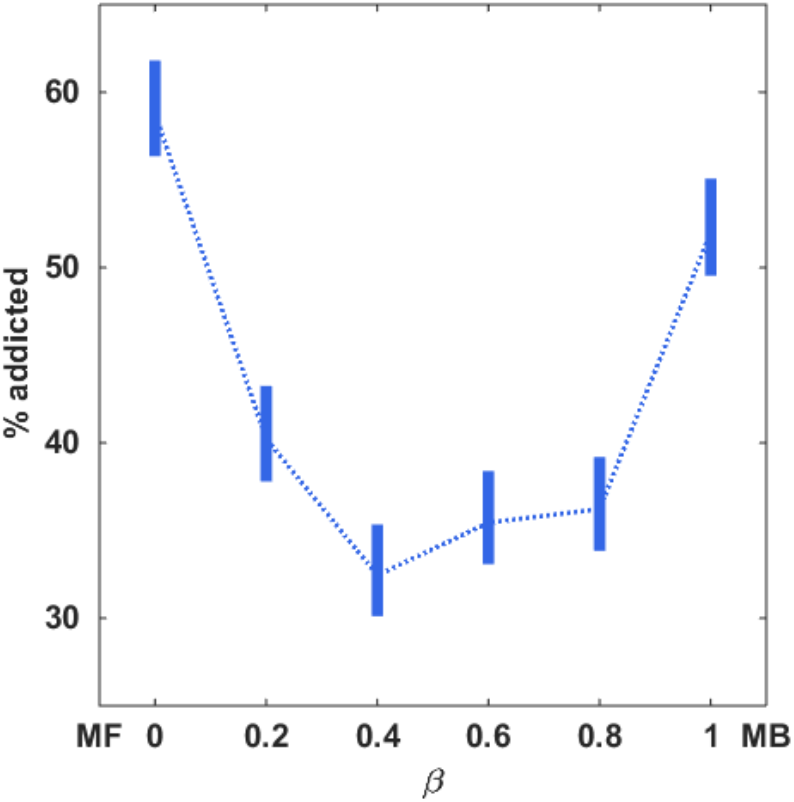
Percentage and confidence intervals of addicted subjects per population, varying *β*. Different *β* values controlling the balance between MF and MB components were used for distinct populations of 900 simulated subjects. Addicted agents are those that during the observation period, 1000 time steps, acquire the addictive reward more often than healthy reward. The percentage of addicted agents per population varied as a function of *β* values, where intermediate values showed a lower percentage of addicted agents (cf. [58]). Confidence interval were estimated assuming two-tail distribution and 95% confidence.

We further investigated these changes in the addiction development probabilities, using the amount of the available cognitive resources as a new independent dimension. The amount of these resources directly determines the depth of navigation in the internal model and, indirectly, how accurately such model is generated. Therefore, limited resources result in incorrect representation and action-sequence assessments, leading to suboptimal choices. To converge to optimum, when the model of the environment is known, the prioritized sweeping algorithm used in the MB requires above 4K updates of the value function. Note that, for these internal *iterations steps*, the value of reaching a state is estimated using the internal model (fixed) without any actual interaction with the world (Table 1). Fig 6 shows that the chances to pursue suboptimal behaviours, i.e. seeking the addictive reward state, are inversely correlated with the resources available for the MB component (which we tested in a range well below the 4K updates necessary for optimal estimation). For instance, the population bounded by 50 Model Based Updates per Step (MBUS) resulted in 50% of subjects expressing addiction-like behaviours after 1K time steps, rising up to 90% of the subjects, after 10K time steps. At the opposite side of the spectrum, populations characterised by high computational resources (e.g. the tested 500 MBUS population) resulted in up to 20% of addicted subjects at 1K time steps, but this percentage falls to 0%, after 10K time steps, showing the agents had developed a correct model of the environment by that moment in the simulation. Contrarily to the MB-MF balance dimension, the behavioural trajectories caused by changes in the available cognitive resources are meaningful only when considered jointly, or in interaction, with the environment complexity. Any increase in the degree of complexity for the environment results in an increased demand of resources, to keep constant the likelihood of convergence to optimum. Ecological environments, however, are not limited by the artificial constraints of a laboratory or simulation set-up, so that they may require prohibitive and biologically implausible amounts of resources and exploration to replicate a result close to the described 500 MBUS population trajectory (see [39, Chapter 8] for related theoretical proofs and [72] for experimental results with state-of-the-art supercomputers over more complex but still simplified environments).

**Figure 6:**
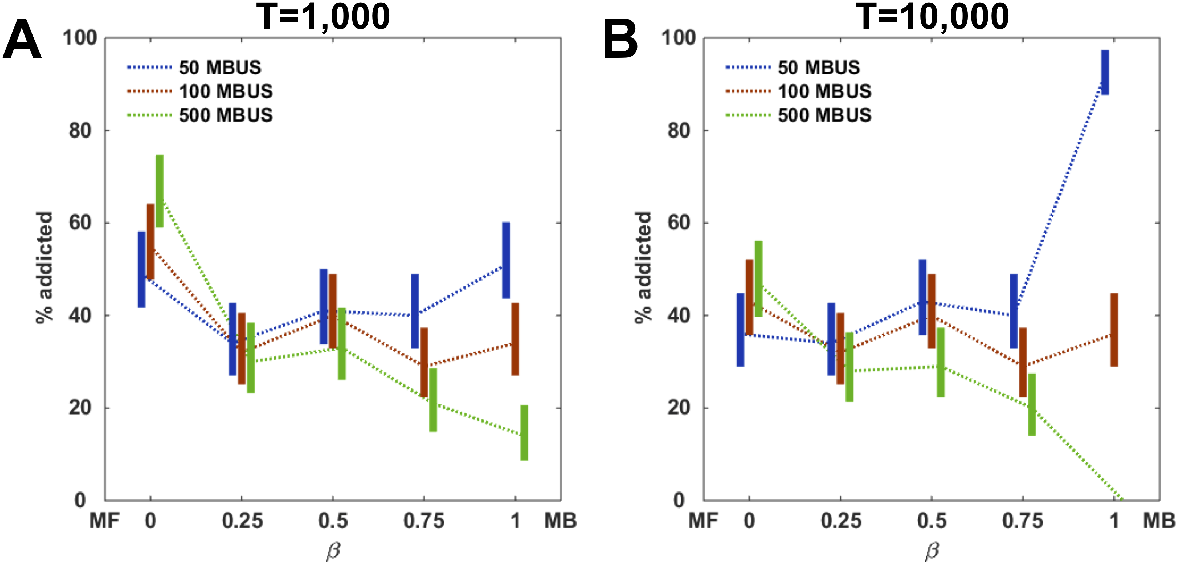
Preference ratios and confidence intervals of agents expressing addiction-like behaviour within each parametrization of cognitive resource bounds (Model Based Updates per Step [MBUS]) and MB/MF balance factor *β*. Initial performance (panel A, analysis on the behaviours in the interval 900 to 1000 timesteps) shows a significant preference for the selection of the addictive reward state, across all values of *β* and most bounds for cognitive resource, with a low for very high resources (500MBUS), in association with *β* = 1. Towards the end of the simulation (panel B, interval 9900 to 10000 timesteps), we found that the populations diverge depending on the amount of cognitive resources available, as preference for the addictive state disappeared in the population characterised by very high resources and *β* = 1. Balanced MB-MF parametrizations (intermediate *β* values) were found generally more resistant to addiction, across values of cognitive bounds. A comparison between panels A and B illustrates the effects of exploration across all the parametrizations. Low values of *β*, dominated by the MF component, slightly reduce the number of addicted subjects after the first 10K steps, for all levels of cognitive resources, as the number of addicted agents remains above one third of the entire population. Exploration and experience with high values of *β* has opposite results, depending on the available cognitive resources. High cognitive resources, jointly with long exploration, lead to a strong reduction of addicted agents, suggesting a correct internal model of the environment is achieved through experience. With low cognitive resources, jointly with a strong MB component (high *β*), experience brings a substantial increase in the number of addicted agents. This result is due to a combination of poor environment representations and limited planning capabilities. Confidence intervals were estimated assuming a two-tail distribution and 95% confidence, with 100 simulated subjects per *β* value.

We hypothesised that the observed behavioural bifurcation, i.e. the diverging behaviours displayed by two identical simulated agents placed in the same environment, was caused by the stochastic nature of the initial exploration phase. We assumed that during this phase, limited knowledge of the environment for both MF and MB components led to non-informative Q-values (i.e. the action-outcome estimations) and therefore to the execution of stochastic action selections. In turn, these initial choices determined which part of the environment would be explored and which would be neglected, shaping the value estimations and further biasing future exploration (cf. [9]).

To test this hypothesis we exposed our agents to the preliminary suboptimal-reward-free simplified environment for a longer time, thus granting early acquisition of an healthy action policy (Fig 7). Under this condition, the agents explored the environment before the introduction of the addictive reward, for a pre-training time (PTT), which lasted a variable number of time steps (50, 200 and 1000). Higher PTT were associated with a better representation of the policy required to reach the healthy state. However, the use of a constant exploration (*ϵ*-greedy) forced the agents to occasionally reach the addictive state reward, after it was introduced in the environment. Despite these exposures to the addictive reward, the chances to develop addiction after a PTT substantially decreased (Fig 7)across values of the parameter *β*, whilst confirming the general resistance to addiction of the balanced MB-MF systems (intermediate values of *β*).

**Figure 7:**
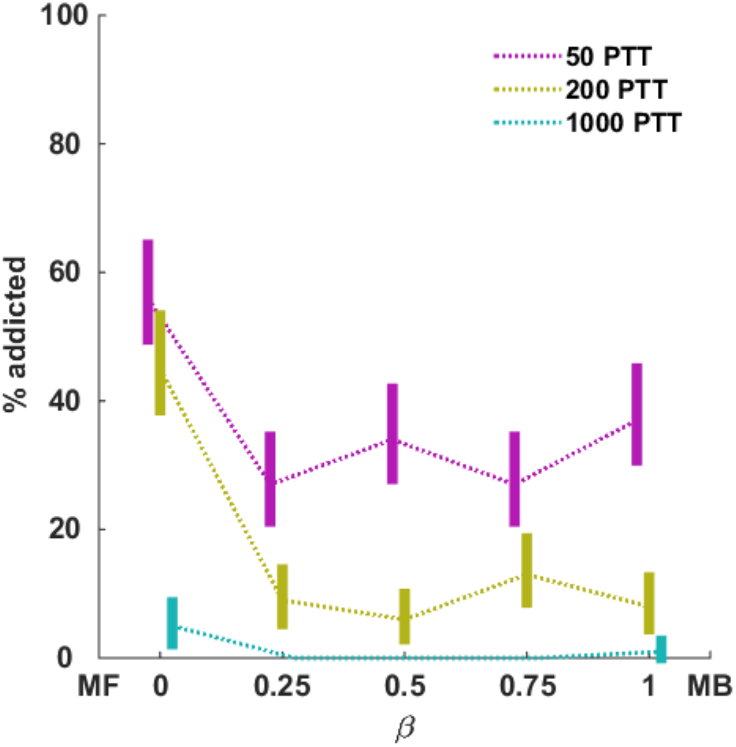
Changes in behavioural trajectories as a function of pre-training time (PTT, timesteps in safe phase) and *β* parameter (MB/MF balance). Exposure to the environment before the introduction of the addictive reward decreased the probability of addiction across all sets of parameters or populations. Extreme values for the parameter regulating MB/MF balance (i.e. *β* ∈ {0,1}) resulted in a residual tendency to addiction even with long exposure. The chart reports confidence intervals for populations tested for 10K steps and composed by 100 agents under each condition, with an evaluation of the behavioural choice selections on the last 1K steps. Confidence intervals were estimated assuming a two-tail distribution and 95% confidence.

Finally, we tested whether sudden environment changes could ignite addiction in agents that had developed the optimal healthy strategy [45, 42]. Our simulations in a Y-maze environment, characterised by the described healthy and addictive reward plus a neutral segment, allowed to test changes in behavioural trajectories after a sudden swap of reward and associated rules between the healthy reward and the neutral segment. This alteration in the environment, taking place after time step 2.5k, when a behavioural policy is consolidated, required the agents to rely again on exploration and learn a new goal directed strategy. The results showed that after this change in the environment, a significant portion of agents previously following a healthy policy developed a sub-optimal addiction behaviour (Fig 8). Importantly, this test proved behavioural shifts to suboptimal behaviour could be induced by changes in the environment, in the absence of malfunctions of the decision components or any pharmacological interference.

**Figure 8:**
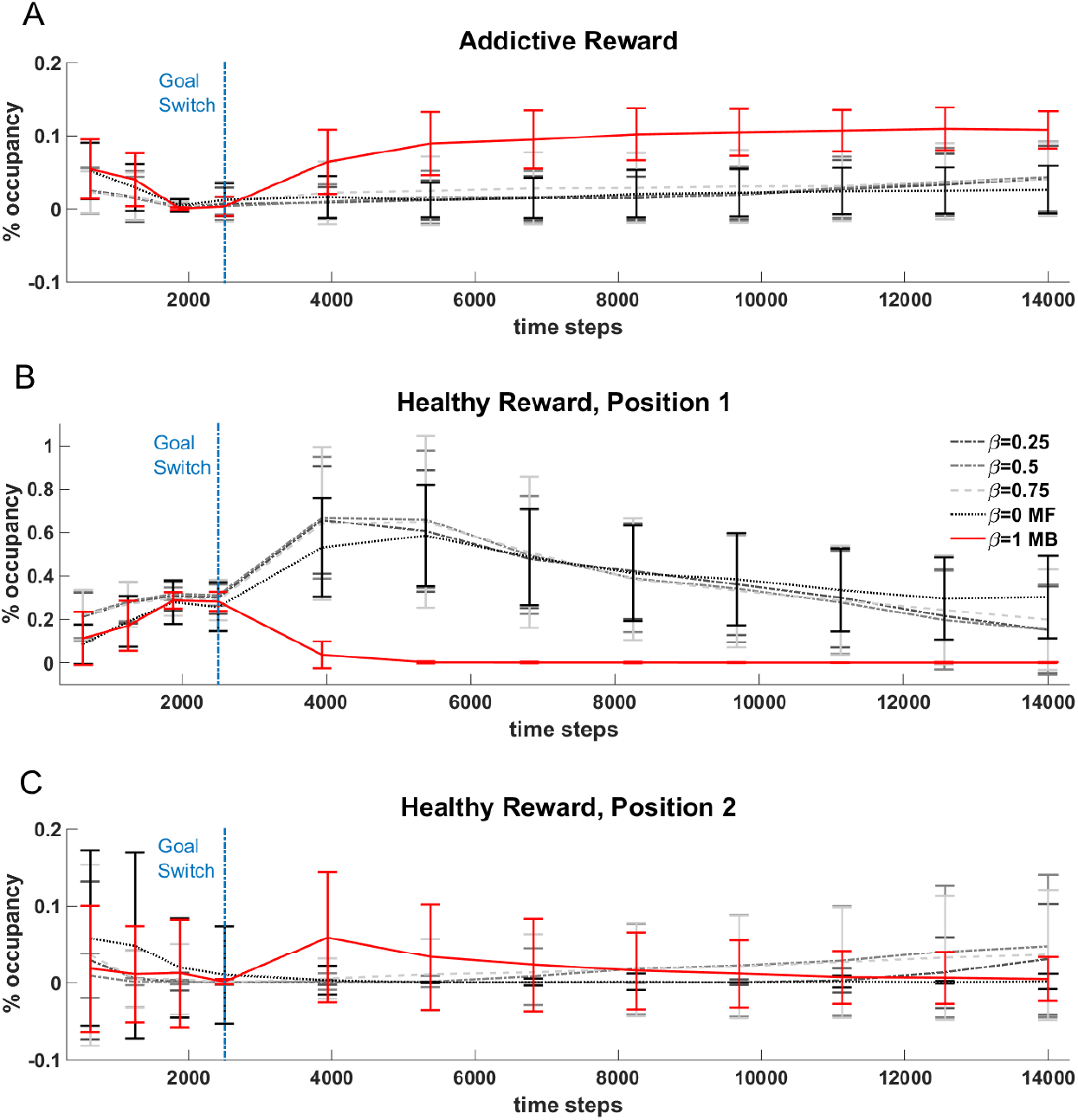
Effects of environment change on healthy subjects. This figure illustrates the effects on behavioural trajectories caused by a change affecting the position of the healthy reward state, depending on the parameter *β* regulating MB-MF balance. The change takes place at time step 2500, when neutral and healthy reward segments are swapped while the addictive segment maintains its configuration. Purely model based agents (*β* = 1) switch rapidly to the addictive behaviour after the change, whereas agents with a non zero MF component gradually unlearn the acquired healthy policy to switch towards either the selection of the addictive state or the re-positioned healthy state. The increased number of visits for the first healthy reward position (panel B) is due to the sudden disappearance of the rewarded action that from this state used to lead (before the swap with the neutral segment) to the starting state (see Fig 2). Without this transition towards the starting state of the environment, the agent expresses cyclic exploratory behaviours, as it can re-enter the now neutral state as soon as it steps outside of it.

## Discussion

As formalised in a seminal work by Redish [16], the RL approach to addiction is based on the hypothesis that drug values are always underestimated by the MF learning system of a biological agent. This phenomenon is mediated by hyper-physiologic phasic responses of dopamine to drug consumption, which deceive the individual consuming the substance of abuse into perceiving the substance itself as always more rewarding than expected (i.e. a non-compensable positive prediction error). In turn, this mismatch between expected and perceived outcomes results in an unlimited growth of the perceived value of drug related actions and aberrant reinforcement, causing habitual decision making, compulsive responses to drug-related stimuli and inelastic behaviour in the face adverse consequences [73, 74, 75, 12].

Despite significant advances in capturing important and complex features of addiction behaviour [19, 18, 11], this model remains primarily an expression of a malfunction of the MF component and therefore it leaves important questions unanswered [76, 20]. In particular, the role of the MB component in addiction is still unclear. First, even though interactions of deliberative computations with dopamine have been described [48], the effects of drug consumption on the generation and assessment of the internal representations of the environment have not been clarified. Second, phenomena such as craving, addiction behaviours which do not rely on stimulus-response habits (e.g. prolonged research for the preferred substance of abuse in novel environments), or non-pharmacological forms of addiction, all seem to suggest that the MB component plays a significant role in driving addiction-like suboptimal behaviours [11].

In this study, we have proposed that addiction-like behaviours can emerge in complex environments, if the dual-learning agent fails to correctly represent and compute action-outcomes associations, due to limited cognitive resources and exploration. In our simulations, a segment of the environment was designed so that an immediate high reward would be followed by multiple, inescapable and heavily stochastic, negative outcomes. We then tested different populations differing in the amount of available cognitive resources and found this variable was inversely correlated with the percentage of agents pursuing the addictive (sub-optimal) reward. Thus, stereotyped inelastic behaviours emerged in a fully accessible and explorable environment, despite the absence of a classic form of drug-induced aberrant prediction error signal or an otherwise malfunctioning MF system. This finding is consistent with previous studies indicating reduced contribution of the MB component may be a risk factor for addiction [77] and we argue it indicates a key computational process underlying those forms of addiction that are not based on the consumption of substances of abuse (e.g. gambling or videogaming).

Beyond the limitations of any experimental settings, the exponential growth of complexity that is associated with ecological environments could easily outmatch the equivalent growth of computational resources in a biologically plausible MB component. Furthermore, our results show that even purely deliberative agents with high cognitive capabilities can still be susceptible to addiction due to dynamic fluctuations in the exploration costs (i.e. sudden changes in the environment), or in the availability of computational resources (e.g. due to stress, a known trigger for addiction [78, 43, 66, 65, 79, 80, 61]). This ambivalency of the MB component in either protecting from or fosetering addiction, depending on the amount of reseources it relies on, is consistent with multiple studies that have highlighted both decreased and increased neural responses in those brain areas associated with MB decision making, in addict individuals in comparison with controls. and depending on task and context [81, 82, 83, 84, 85].

This MB vulnerability can interact with previously described ones [3]. In forms of addiction dominated by the non-compensable prediction errors and hyper-physiologic DA responses, erroneous representations and assessments of the environment can aggravate the behavioural symptoms associated with the classic MF malfunction. This interaction can account for those complex non-habitual drug-seeking behaviours that are not triggered by the presence of drug-related stimuli [23, 18, 3, 24]. Importantly, a resource bounded MB component may fail in evaluating long term action effects even after extensive exploration, so that even after the MF component has eventually converged towards an optimal behaviour (e.g. after a successful treatment), the MB component may keep pursuing sub-optimal policies, contributing to both craving and relapse [86]. Furthermore, by over-selecting the addictive reward early on in the task, exploration and representation of alternative routes in addicted agents remain limited, so that the stronger the addiction, the more compromised the model of the environment. This phenomenon, jointly with the fluctuations of long term outcome estimations under conditions of low MB resources [39, Sections 2.[4–5], results in lowering the chances to disengage from pursuing the suboptimal policy at each step taken in the direction of the addictive reward, putatively simulating a context-related sense of inability to stop [19].

Finally, the vulnerability we have described can be seen as ideally contiguous with those associated with state identification errors [9, 87, 88, 89, 90]. Under conditions of the environment in which information about the states is either incomplete or inaccessible, the resulting interaction between state identification and value estimation can cause the creation of fictitious internal states, where addictive behaviours would always be considered as highly rewarding [9]. This hypothesis was originally proposed as a cause of context-driven addiction and has been used to describe gambling [9]. Under the conditions we have proposed, information exceeding an agent cognitive capabilities would be essentially lost to an agent, however the two vulnerabilities remain significantly different under many other aspects. The vulnerability we have described is not restricted to the opacity of a specific environment, and the dynamic interplay between exploration demands and availability of resources allowed us to account for the presence of different behavioural trajectories or phenotypes. We have observed that behavioural differences can arise from any change (either temporary or permanent) in the key parameter of the available cognitive resources, as well as unexpected changes in the environment structure or simply due to less than *few hundreds* initial stochastic exploration steps. These differentiations and behavioural trajectories took place despite the presence of a converging MF algorithm (as demonstrated in the *long run* tests) and it was neither caused by a disruption of the classical TD-MF learning mechanism [16, 19], nor by incomplete access to information concerning rewards and punishments in the environment [9].

Our findings have interesting implications for treatment development. A crucial problem is that the MB component is unlikely to increase its computational power with training, so that even if a correct model is formed, the agent might still pursue addictive behaviours, initiating relapse, due to difficulties in assessing complex ramifications associated with apparently rewarding initial choices. Thus, we hypothesise a treatment could aim at simplifying or making more explicit and accessible the structure of the environment. In doing so, normally occurring negative outcomes associated with the addictive behaviour would be easier to be taken into consideration and-importantly-courses of action leading to healthy policies would become competitive in the MB component. Unfortunately, there is the possibility that, independent of treatment, the MB component might keep associating a high reward to the addictive behaviour due to a stochastic representation of past experienced rewards, possibly modulated by reward intensity and distance in time. We hypothesise these conditions could be ameliorated by a conflict between MF and MB component, where addiction-avoiding habits could be developed during treatment, as suggested by our pre-training tests (Fig 7).

In conclusion, several studies focus on the effects that different sources of complexity (most prominently, social factors [91, 92] and stress [93, 45]) may have on addiction, however current computational modelling literature has often neglected these aspects [29, 31]. In this work we have proposed a step forward in the direction of more ecologically plausible simulations of healthy and dysfunctional behaviours, as we highlighted the interaction between limited MB resources and overwhelming representation requirements.

